# Converging structural and functional connectivity of orbitofrontal, dorsolateral prefrontal, and posterior parietal cortex in the human striatum

**DOI:** 10.1101/006619

**Authors:** Kevin Jarbo, Timothy D. Verstynen

**Affiliations:** Department of Psychology, Center for the Neural Basis of Cognition Carnegie Mellon University, Pittsburgh, PA

## Abstract

Modification of spatial attention via reinforcement learning (Lee & Shomstein, 2013) requires the integration of reward, attention, and executive processes. Corticostriatal pathways are an ideal neural substrate for this integration because these projections exhibit a globally parallel (Alexander, De Long, and Strick, 1985), but locally overlapping (Haber, 2003), topographical organization. Here, we explored whether there are unique striatal regions that exhibit convergent anatomical connections from orbitofrontal cortex (OFC), dorsolateral prefrontal cortex (DLPFC), and posterior parietal cortex. Deterministic fiber tractography on diffusion spectrum imaging data from neurologically healthy adults (N=60) was used to map fronto-and parieto-striatal projections. In general, projections from cortex were organized in a rostral-caudal gradient along the striatal nuclei; however, we also identified two bilateral convergence zones—one in the caudate nucleus and another in the putamen—that consisted of voxels with projections from OFC, DLPFC, and parietal regions. The distributed cortical connectivity of these striatal convergence zones was confirmed with follow-up functional connectivity analysis from resting state fMRI data from 55 of the participants, in which a high percentage (62-80%) of structurally connected voxels also showed significant functional connectivity. These results delineate a neurologically plausible network of converging corticostriatal projections that may support the integration of reward, executive control, and spatial attention that occurs during spatial reinforcement learning.

## Introduction

Contextual factors, including cue/target proximity and their co-occurrence within the same bounded object, can bias visuospatial attention (Egeth & Yantis, 1997; Posner, Snyder, & Davidson, 1980). However, recent research has shown that high reward targets can override these spatial- and object-based attentional biases (Lee & Shomstein, 2013a). Reinforcement learning (Sutton & Barto, 1998) is traditionally described as a process of incorporating feedback from previous trials to maximize the likelihood of reward on future action selections. Together, these findings suggest that reinforcement learning can alter bottom-up influences of stimulus features on attentional allocation.

Functionally, reinforcement learning depends on the striatum (Daw & Doya, 2006; Dayan & Abbott, 2001; O’Doherty, 2004; Knutson, Westdorp, Kaiser, & Hommer, 2000). Most studies focus on reward processing by the ventral striatum at the local level (O’Doherty, Dayan, Friston, Critchley, & Dolan, 2003; Mcclure, Berns, & Montague, 2003; Mcclure, York, & Montague, 2004; Pagnoni, Zink, Montague, & Berns, 2002; Rodriguez, Aron, & Poldrack, 2006), but evidence of dorsomedial caudate involvement in reward-based response updating (Delgado, Locke, Stenger, & Fiez, 2003; Delgado, Miller, Inati, & Phelps, 2005; Knutson & Cooper, 2005; Kuhnen & Knutson, 2005; Lohrenz, McCabe, Camerer, & Montague, 2007) suggests a more global role of the striatum in behavioral updating. While the striatum is generally viewed as a central integration point of cortical information via strictly parallel circuits (Alexander, DeLong, & Strick, 1986), there is evidence for overlap from spatially disparate cortical areas (Haber, 2003). Through this diffuse overlap of corticostriatal projections, the striatum may provide a substrate for reinforcement learning that integrates reward and executive control signals, respectively, from orbitofrontal (OFC) and dorsolateral prefrontal cortex (DLPFC; see Haber & Knutson, 2010 for review).

Evidence of reinforcement learning effects on spatial attention comes from studies showing improved visual search proficiency in high-reward locations over low-reward locations (Della Libera & Chelazzi, 2006; Kristjansson, Sigurjonsdottir, & Driver, 2010; Lee & Shomstein, 2013a, 2014). This form of spatial attention is functionally associated with posterior parietal cortex in humans and nonhuman primates (Colby & Goldberg, 1999; Silver, Ress, & Heeger, 2005). Nonhuman primate histology research has shown a pattern of parietostriatal connectivity where posterior parietal projections terminate in distributed clusters along the caudate nucleus, proximal to OFC and DLPFC projection termination sites (Cavada & Goldman-Rakic, 1991; Selemon & Goldman-Rakic, 1985, 1988), that has also been confirmed functionally in humans (Choi, Yeo, & Buckner, 2012; Di Martino *et al*., 2008). Taken together, this evidence implicates the striatum as a potential integration point for reward, executive control, and spatial attention; however, the specific pattern of convergent inputs from the associated cortical areas has not been confirmed.

Using diffusion spectrum imaging (DSI) and resting state fMRI, we explore a neurologically plausible network of converging white matter projections in the striatum that may support the integration of information from OFC, DLPFC, and posterior parietal cortex. The observation of overlapping corticostriatal projections and corresponding functional connectivity would provide evidence for a structurally and functionally integrative network that may underlie mechanisms of spatial reinforcement learning.

## Materials & Methods

### Participants

Sixty participants (28 male, 32 female) were recruited locally from the Pittsburgh, Pennsylvania area as well as the Army Research Laboratory in Aberdeen, Maryland. Participants were neurologically healthy adults with no history of head trauma, neurological or psychological pathology. At the time of data acquisition, participant ages ranged from 18 to 45 years old (mean age 26.5 years old). Informed consent, approved by the Institutional Review Board at Carnegie Mellon University, also in compliance with the Declaration of Helsinki, was obtained for all participants. Participants were all financially compensated for their time.

### MRI Acquisition

All 60 participants were scanned at the Scientific Imaging and Brain Research (SIBR) Center at Carnegie Mellon University on a Siemens Verio 3T magnet fitted with a 32-channel head coil. A magnetization prepared rapid gradient echo imaging (MPRAGE) sequence was used to acquire a high-resolution (1mm^3^ isotropic voxels, 176 slices) T1-weighted brain image for all participants. Diffusion spectrum imaging (DSI) data was acquired following fMRI sequences using a 50-minute, 257-direction, twice-refocused spin-echo EPI sequence with multiple q values (TR = 11,400ms, TE = 128ms, voxel size = 2.4mm^3^, field of view = 231 × 231mm, b-max = 5,000s/mm^2^, 51 slices). Resting state fMRI (rsfMRI) data consisting of 210 T2*-weighted volumes were collected for each participant with a blood oxygenation level dependent (BOLD) contrast with echo planar imaging (EPI) sequence (TR = 2000ms, TE = 29ms, voxel size = 3.5mm^3^, field of view = 224 × 224mm, flip angle = 79°). Head motion was minimized during image acquisition with a custom foam padding setup designed to minimize the variance of head motion along the pitch and yaw rotation directions. The setup also included a chin restraint that held the participant’s head to the receiving coil itself. Preliminary inspection of EPI images at the imaging center showed that the setup minimized resting head motion to about 1mm maximum deviation for most subjects.

### Diffusion MRI Reconstruction

DSI Studio (http://dsi-studio.labsolver.org) was used to process all DSI images using a q-space diffeomorphic reconstruction method (Yeh & Tseng, 2011). A non-linear spatial normalization approach (Ashburner & Friston, 1999) was implemented through 16 iterations to obtain the spatial mapping function of quantitative anisotropy (QA) values from individual subject diffusion space to the FMRIB 1mm fractional anisotropy (FA) atlas template. The orientation distribution functions (ODFs) were reconstructed to a spatial resolution of 2mm^3^ with a diffusion sampling length ratio of 1.25. Whole-brain ODF maps of all 60 subjects were averaged to generate a template image of the average tractography space.

### Fiber Tractography & Analysis

A September 23, 2013 build of DSI Studio was used to perform all tractography by implementing an ODF-streamline version of the FACT algorithm (Yeh, Verstynen, Wang, Fernández-Miranda, & Tseng, 2013). Region-of-interest (ROI)-based tractography was carried out to isolate streamlines between pairs of ipsilateral ROI masks selected from the SRI24 Multi-Channel atlas (Rohlfing, Zahr, Sullivan, & Pfefferbaum, 2010). Caudate nucleus and putamen masks were merged to generate a striatum mask that was then expanded by one voxel (2mm) in all directions. Twenty-six cortical surface masks (13 per hemisphere) in the frontal and parietal lobes were selected from the SRI24 atlas as targets for corticostriatal tractography, including: gyrus rectus (Rectus); ventromedial prefrontal cortex (Frontal_Med_Orb); opercular, orbital and triangular parts of the inferior frontal gyrus (Frontal_Inf_Oper, Frontal_Inf_Orb, Frontal_Inf_Tri); dorsal and orbital middle and superior frontal gyri (Frontal_Mid, Frontal_Mid_Orb, Frontal_Sup, Frontal_Sup_Orb); superior and inferior parietal lobules (Parietal_Sup, Parietal_Inf); angular gyrus (Angular) and supramarginal gyrus (SupraMarginal).

Fiber tractography (Figure 1) was initiated from seed positions with random locations within the seed mask with random initial fiber orientations. Using a step size of 1mm, the directional estimates of fiber progression within each voxel was weighted by 80% of the incoming fiber direction and 20% of the previous moving direction. A streamline was terminated when the QA index fell below 0.05 or had a turning angle greater than 75°. The QA threshold was set to 0.04 for tracking streamlines from the dorsal middle frontal gyri (Frontal_Mid) due to detection of significantly fewer corticostriatal projections than expected (Verstynen, Badre, Jarbo, & Schneider, 2012). Each cortical surface ROI mask was paired with an ipsilateral striatum ROI mask, both of which were designated as ends in DSI Studio, and whole-brain seeded tractography continued for 3×10^8^ seeds (approximately 3000 samples per voxel in the whole brain mask). To be included in the final dataset, streamlines had to 1) have a length less than 120mm, and 2) terminate in the cortical surface mask at one end and within the ipsilateral striatum mask at the other. To facilitate further analyses, the 13 sets of streamlines for every pairing in each hemisphere were combined into four meta-regions. The OFC meta-region was comprised of streamlines from medial and lateral OFC, including: gyrus rectus (Rectus), the orbital part of the inferior frontal gyrus (Frontal_Inf_Orb) and middle (Frontal_Mid_Orb) and superior frontal (Frontal_Sup_Orb) gyri. The DLPFC meta-region consisted of streamlines from opercular (Frontal_Inf_Oper) and triangular (Frontal_Inf_Tri) parts of the inferior frontal gyrus, as well as middle (Frontal_Mid) and superior frontal (Frontal_Sup) gyri. Streamlines from the superior (Parietal_Sup) and inferior parietal lobules (Parietal_Inf), angular gyrus (Angular), and supramarginal gyrus (SupraMarginal) constituted the parietal meta-region. For a more complete assessment of the endpoint distributions of these meta-regions, streamlines from ventromedial prefrontal cortex (vmPFC; Frontal_Med_Orb) tractography were included.

**Figure 1.**
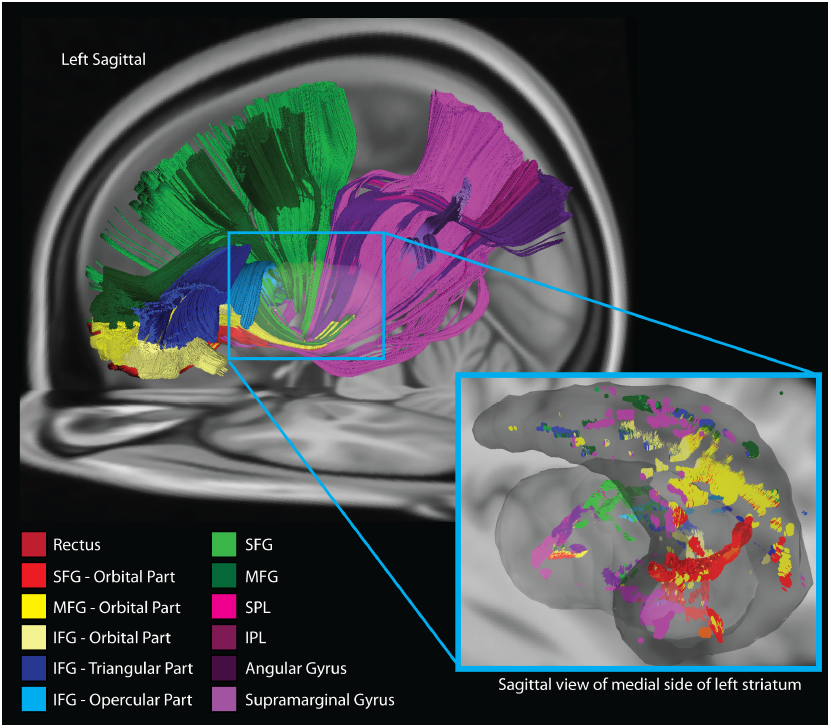
Tractography analysis on the average subject template brain (CMU-60 template). Each pathway was tracked using a region of interest approach between a specific cortical area and a mask of the striatal nuclei (gray region in inset). Red and yellow streamlines represent medial and lateral orbitofrontal cortex (mOFC and lOFC), respectively. Green and blue represent dorsolateral prefrontal cortex (DLPFC), while violet streamlines indicate projections between parietal cortical regions and the striatum. Inset image shows a medial view of the streamline endpoints from each cortical pathway.

Upon completion of tractography, custom MATLAB functions were used to generate four striatal endpoint density maps (i.e., convergence zones, see Figure 4A) where all cortical meta-regions yielded overlapping projections within ipsilateral striatum. First, the three-dimensional coordinates of the streamline endpoints in the caudate nucleus and putamen within each hemisphere were extracted. To obtain matrices of striatal endpoint coordinates for each meta-region for all participants, a mask for each caudate nucleus and putamen were loaded separately into MATLAB with streamlines from each ipsilateral cortical region. Striatal endpoints were extracted and saved as a new NIfTI mask, resulting in a convergence zone representing the total volume of contiguous voxels (cluster size k > 20) within each nucleus where termination points of projections from the OFC, DLPFC and parietal meta-regions were detected. This was done for both caudate nuclei and putamen resulting in four (left caudate, left putamen, right caudate, right putamen) convergence zone masks for all 60 diffusion imaging datasets. Next, convergence zone masks for each nucleus were used to calculate maps of the mean convergence zone as well as to assess the consistency and significance of convergence zone volumes across all datasets. The significance at each convergence zone was calculated using a one-sample t-test across all subjects with an FDR-corrected threshold (q) of less than 0.05.

To confirm the pattern of connectivity observed through the constrained, ROI-based approach, tractography (Figure 4B) was reseeded from a whole-brain mask with each convergence zone designated as an end. This was repeated separately for all four convergence zone masks in all 60 datasets. Tractography was initiated using the same parameters described in the *Fiber Tractography* section with the only differences being that all fibers were generated using a QA of 0.05, and tracking proceeded until 50,000 fibers were detected, rather than 3×10^8^ seeds. The three-dimensional endpoints resulting from reseeded tracking for each convergence zone were extracted to obtain matrices of the endpoints that were then converted into NIfTI-format density maps for all 60 datasets. Mean maps of cortical endpoint terminations for each of the four convergence zones were calculated, resulting in an averaged density of endpoints throughout the brain. Maps of t and q values from a one-sample t-test were also calculated from these data to evaluate the consistency and significance of endpoint terminations from the reseeded tractography across all datasets.

### Striatal Endpoint Distribution Analysis

In order to capture the distribution of endpoints from fibers originating in the meta-regions-of-interest masks for the OFC, DLPFC, and parietal lobes, we calculated the probability of observing a fiber along the sagittal (i.e., the y-coordinate) plane for each subject. For this analysis, the y location in MNI-space of each streamline endpoint within the striatum was extracted. This position vector was binned at 0.5mm increments and the number of endpoints detected in each bin was isolated. This was then converted to a probability function by dividing the number of endpoints detected in each bin by the total number of streamlines terminating within the striatum mask. This analysis was performed separately for the meta-region cortical streamlines terminating in the caudate and putamen masks.

### Resting State fMRI Preprocessing and Analyses

SPM8 (Wellcome Department of Imaging Neuroscience, London, UK) was used to preprocess all resting state fMRI data (rsfMRI) collected from 55 of the 60 participants with DSI data. The transformation to normalize each EPI dataset was carried out by first weighting each individual EPI (i.e., source) image with a mean EPI image constructed from all 55 datasets. An EPI template reconstructed in MNI-space, supplied with SPM, was used as a target for estimating the normalization of the individual source images. Then, the smoothing kernel was set to a FWHM of 4mm and the SPM8 default parameters were used for all other estimation options to generate a transformation matrix to be applied to the source image. Finally, the transformation matrix was applied to each volume of the individual source images weighted by the group mean EPI for further analyses.

The convergence zones obtained from the tractography analyses were used as seed points for the functional connectivity analysis. A series of custom MATLAB functions were used to 1) extract the voxel time series of activation for each convergence zone, 2) remove noise from white matter and cerebrospinal fluid (CSF) time series, 3) smooth the de-noised time series data, and 4) calculate t and p values of consistent activation with corresponding significance. Resting state fMRI data was analyzed using AFNI (Cox, 1996) to calculate functional activity throughout the brain correlated with each convergence zone seed in accordance with previously employed methods (see Choi *et al*., 2012). Specifically, functional activity correlations (r) were converted to *Z-*scores using Fisher’s r-to-*Z* transformation for each convergence zone across all 55 datasets.

First, a convergence zone was loaded into MATLAB 8.1/R2013a (The Mathworks, Sherborn, MA) with an individual subject’s rsfMRI time series data. The time series of activation corresponding with the volume of the convergence zone mask was extracted, yielding activation values for each voxel in the mask across all 210 volumes of the rsfMRI BOLD EPI sequence. Next, the convergence zone time series was de-noised by regressing the time series information of manually selected white matter and CSF voxels out of the total convergence zone time series activation. Once de-noised, the data were smoothed with a Gaussian kernel (FWHM = 2mm) and a one-sample t-test was run to identify consistent, significant functional activity correlated with the convergence zone time series across all 55 individual datasets (Figure 5). Corresponding FDR-corrected q values were also calculated for each convergence zone and visualized as Z-maps initially thresholded at q < 0.05 for significance at Z = 1.96.

### Structural and Functional Connectivity Overlap Analysis

Using a custom MATLAB function, t-maps of consistent structural connectivity from the DSI data, and *Z*-transformed correlation (r) maps from the fMRI data were used to calculate the percentage of structurally significant voxels (i.e., a cortical voxel that had significant structural connectivity with a striatal convergence zone) that were also functionally significant. For this, the DSI t-map data were thresholded at q < 0.05 to yield all significant voxels with structural connections that were consistent across all 60 DSI datasets. Corresponding rsfMRI data were also thresholded at q < 0.05, resulting in maps of voxels with significant functional connectivity across all 55 fMRI datasets. For each convergence zone, t-maps and *Z-*maps of structural and functional connectivity, respectively, were loaded into MATLAB. A voxel was considered to have significant structural or functional connectivity if the one-sample t-test to find consistent connections across all DSI or rsfMRI datasets resulted in a significant *q-*value. The maps of significant structural and functional connectivity for each convergence zone were binarized such that all voxels with a q < 0.05 were set to 1, and all other voxels were set to 0. After transforming the binary data into single column vectors, the dot product of significant structural and functional voxels was summed and divided by the number of significant structural voxels. This calculation yielded the percentage of cortical voxels that had significant structural and functional connectivity with a striatal convergence zone, aggregated across all voxels within a given zone. A permutation test was conducted to determine chance levels of overlap between the structural and functional measures. On each iteration of the permutation test, the vector of significant functional voxels was randomly permuted and the percent overlap with the structural vector recalculated. This process was repeated for 1000 iterations for each convergence zone ROI to construct the 95% confidence interval of chance overlap between structural and functional overlap (i.e., to construct the null distribution that structurally connected voxels to the convergence zone randomly overlapped with functionally connected voxels).

## Results

### Topography of corticostriatal projections

Across the 13 cortical regions evaluated, we observed a consistent topography of striatal endpoint fields that was in agreement with previously reported patterns found in both humans (Draganski *et al*., 2008; Lehéricy, Ducros, Van de Moortele, *et al*., 2004; Verstynen *et al*., 2012) and non-human primates (Cavada & Goldman-Rakic, 1991; Haber, Kunishio, Mizobuchi, & Lynd-Balta, 1995; Haber, Kim, Mailly, & Calzavara, 2006; Haber & Knutson, 2010; Selemon & Goldman-Rakic, 1985, 1988). Figure 1 shows a sagittal view of the frontal and parietal projections into the ipsilateral striatum from a template brain, reflecting the average voxel-wise diffusion information across all participants. As expected, streamlines were grouped together spatially according to their cortical origin, with orbitofrontal and dorsolateral prefrontal streamlines bundled in more anterior regions than streamlines from parietal cortex. This is most clearly evident in the inset panel of Figure 1, showing the corresponding endpoints from each set of streamlines within the caudate nucleus (dark gray) and putamen (light gray) of the left striatum. In particular, ventromedial PFC and medial OFC (vmPFC and mOFC; red) and lateral OFC (lOFC; yellow) regions both show dense clustering within the rostral and ventromedial striatum. DLPFC (blue and green) clusters were observed dorsal and lateral to OFC clusters. Finally, parietal (violet) clusters were widely distributed throughout the striatum and observed posterior to both frontal clusters, consistent with previous histological work (Selemon & Goldman-Rakic, 1985; Cavada & Goldman-Rakic, 1991). Though OFC, DLPFC and parietal projections show a rich, distinct topography in the striatum, a marked overlap of their termination fields is observed in both the caudate nuclei and putamen, bilaterally.

In order to determine the consistency of this topographic organization across individuals, we then looked at the common regions of endpoint densities in the striatum for all 60 participants. For ease of analysis, all 13 ROIs were collapsed into four meta-regions: ventromedial PFC (vmPFC), orbitofrontal cortex (OFC), dorsolateral prefrontal cortex (DLPFC) and parietal (see Methods for more details). Figure 2 shows these endpoint fields for each meta-region cluster.

**Figure 2.**
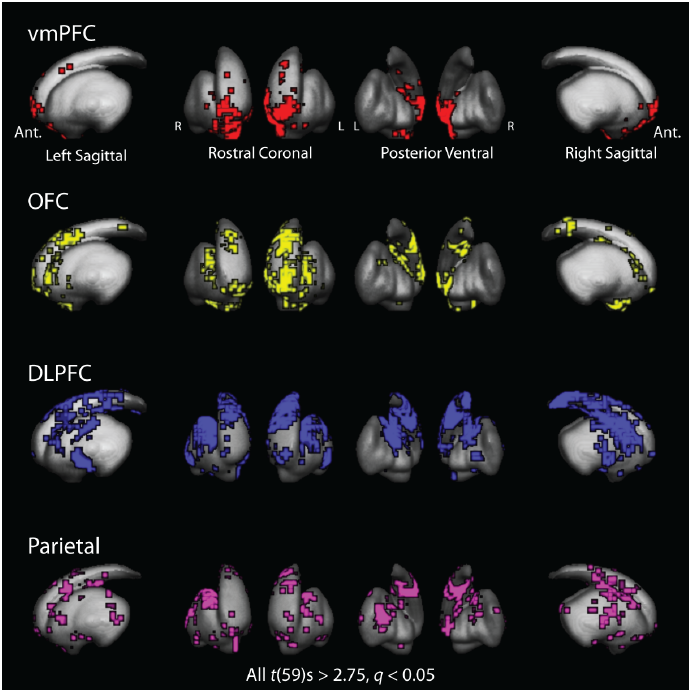
Group statistical maps of common endpoint locations from four cortical meta-regions: ventromedial prefrontal cortex (vmPFC; red), orbitofrontal cortex (OFC; yellow), dorsolateral prefrontal cortex (DLPFC; blue) and parietal cortex (violet). Voxels indicate regions with significant endpoint densities from cortex determined using a 1-sample t-test and corrected for multiple comparisons.

As expected, the endpoint clusters of projections from the four meta-regions exhibited similar topographical distributions. At this group level, a distinct rostral-to-caudal shift was present in endpoint distributions with more anterior cortical surface regions having a higher concentration of endpoints rostrally within the striatum, while projection endpoints from more posterior cortical regions were concentrated caudally (Figure 2). Specifically, vmPFC (red) projections terminated in the most rostral and ventromedial striatal regions, followed caudally and dorsally by OFC (yellow), DLPFC (blue) and parietal cortex (violet). This rich, topographical organization of cortical projection endpoints along the striatum demarcates a distinct spatial segmentation of cortical inputs, while also providing evidence of some local overlap of corticostriatal projections from adjacent cortical networks.

### Convergence of corticostriatal projections

As shown in Figure 2, the primary topographical shift of corticostriatal endpoint distributions is in the rostral-caudal direction. Indeed, previous work has suggested that the sagittal plane is the primary gradient of projection overlap in the striatum (Draganski *et al*., 2008; Selemon & Goldman-Rakic, 1985, 1988; Verstynen *et al*., 2012). In order to see if the major projection fields also exhibit a degree of overlap along this plane, we examined the distribution of fiber streamline endpoints on the striatum along the sagittal (i.e., y) axis (Figure 3). Overall, the DLPFC and OFC peak densities are similar across both hemispheres, with the OFC showing an extra peak in the probability density function (PDF) in the anterior aspects of the striatum, likely reflecting projections into the shell of the caudate. In contrast, the highest peaks for the parietal cortex were shifted in a consistently caudal direction compared to both the DLPFC and OFC (Fig. 3C). While there is a discernable shift in the peaks of the PDFs from these three cortical areas, there is also substantial overlap along the sagittal plane. In particular, the parietal PDF extends in a rostral direction, into the regions of highest density for OFC and DLPFC projections.

**Figure 3.**
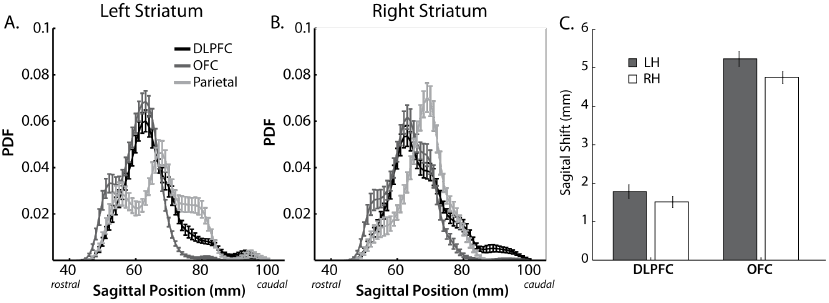
Probability density function of streamline endpoint locations along the sagittal plane of the caudate nucleus. A, B) Black PDF corresponds with DLPFC projections, dark gray PDF with OFC, and light gray PDF with parietal cortex. The horizontal axis represents the rostral-to-caudal location of the peak densities along the sagittal axis with the values of the PDFs represented on the vertical axis of the graph. C) Peak shift between the parietal PDF and each prefrontal PDF across all subjects. Vertical axis shows the difference as parietal minus frontal, with positive values indicating a caudal shift in millimeters. Error bars show standard error of the mean.

To see if this area of overlapping densities reflects convergent projections, we used a conjunction analysis to identify voxels that had significant endpoint densities from OFC, DLPFC, and parietal masks (see Materials and Methods). Clusters of these conjunction voxels (k > 20) were isolated bilaterally within the caudate nucleus and putamen separately. Each nucleus contained a distinct cluster of these convergent fields that appeared to be relatively symmetric across hemispheres (Figure 4A). In the caudate, the convergence zones were isolated along the rostral portion of the body of the caudate. In the putamen, the convergence zones were found on the dorsal and rostral aspects of the nucleus. It is important to note that these clusters are away from ventral striatal regions that are typically thought of as the main termini of OFC projections (Haber, 2003).

**Figure 4.**
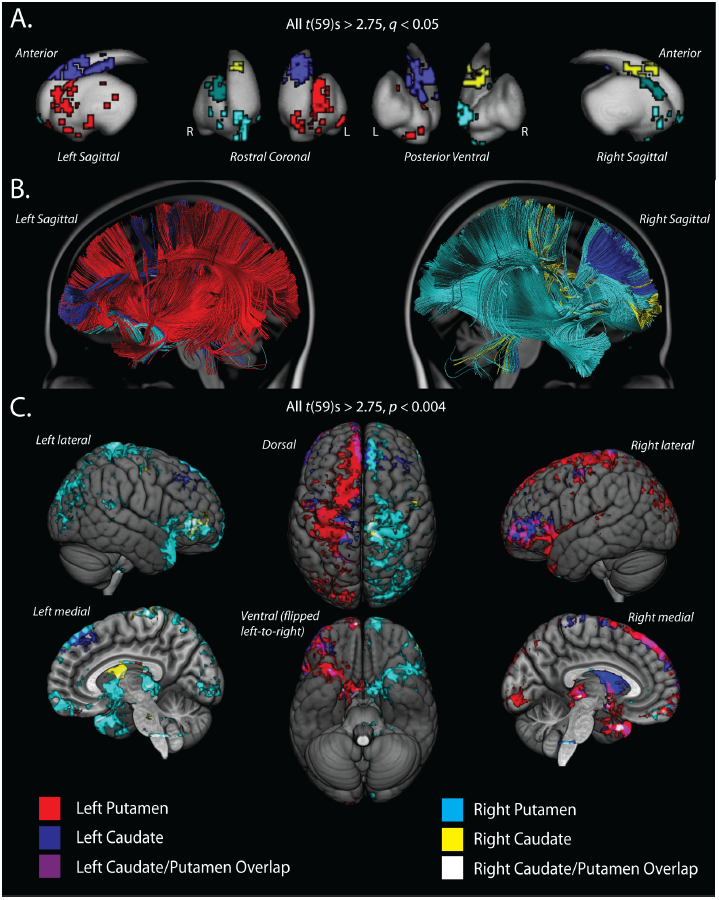
A) Convergence zones of OFC, DLPFC and Parietal projections along the striatal nuclei. These were identified using a conjunction analysis of significant voxels containing projections from all three cortical meta-regions and segmented using k-means clustering. Two convergence zones on each nucleus were identified: left putamen (red), left caudate (blue), right putamen (cyan), right caudate (yellow). A cluster threshold of k > 20 was used and voxels that appear isolated on the surface of the striatum are indeed connected to a larger, contiguous cluster of voxels deeper within the nuclei. B) Tractography on the CMU-60 template brain into each of the convergence zone masks. Unlike the tractography shown in Figure 1, cortical endpoints were not constrained by *a priori* region masks. Streamline colors match scheme shown in *A*. C) Significant endpoint locations, across subjects, for each striatal convergence zone. Colored voxels show regions that had consistent endpoint projections into a convergence zone, based on a 1-sample t-test (uncorrected) across all subjects. Connections ending in the putamen convergence zone originate from a much larger and more distributed set of cortical areas than the streamlines that end in the caudate convergence zone.

In order to get a more complete picture of cortical projections into the striatal convergence zones, we performed a second whole-brain tractography analysis, isolating only streamlines that ended in each of the clusters shown in Figure 4A. Figure 4B shows streamlines that end in the convergence zones from tractography performed on the CMU-60 template. Generally, both putamen convergence zones showed more distributed projections (Figure 4B: left, red; right, cyan) than the caudate convergence zones projections (Figure 4B: left, blue; right, yellow). This pattern held when we looked at the consistent cortical endpoint locations across all 60 participants (Figure 4C; all t(59)s > 2.75, uncorrected p < 0.004). A high proportion of connections between OFC, DLPFC, sensorimotor, parietal, occipital and posterior temporal cortex and the putamen are observed in both hemispheres. Additionally, there is a substantial overlap of ipsilateral projections into both caudate nuclei and putamen from lateral OFC, anterior DLPFC, primary motor and sensorimotor areas indicated by purple and white regions in the left and right hemispheres, respectively. In general, the connectivity with the putamen was much more distributed across the frontal and parietal regions than the caudate connectivity. In frontal cortex, the caudate connectivity was isolated to regions of the lateral OFC, precentral gyrus and small sections of the medial wall of the superior frontal gyrus. Within the parietal cortex, caudate connectivity was restricted to the very anterior portion of the intraparietal sulcus and portions of the postcentral gyrus and sulcus, near somatosensory regions. Interestingly, since our original tractography was restricted to ipsilateral corticostriatal projections, left caudate tracking resulted in notable intra- and interhemispheric connections with presupplementary motor areas including the dorsal and medial superior frontal gyrus as well as the lateral middle frontal gyrus areas. The observed contralateral connectivity between left caudate convergence zone and right dorsolateral prefrontal areas is indeed consistent with nonhuman primate histology (McGuire, Bates, & Goldman-Rakic, 1991) and human diffusion imaging work (Lehéricy, Ducros, Krainik, *et al*., 2004). However, we did not observe consistent projections between the right caudate convergence zones and ipsilateral or contralateral presupplementary motor areas. Putamen connectivity to frontal regions, however, was distributed across broad swaths of cortex including medial and lateral OFC, medial and lateral DLPFC and the precentral gyrus. In parietal regions, the putamen was connected to most of the dorsal posterior parietal cortex, including posterior regions of the intraparietal sulcus, angular gyrus and parietal occipital border. While these differences between the striatal nuclei connectivity implicate a more distributed cortical connectivity to the putamen convergence zone than the caudate convergence zone, it is important to point out that structural connectivity from the OFC, DLPFC, and parietal meta-regions are consistently observed across all datasets, confirming our original region-of-interest analysis.

### Functional Connectivity of Convergence Zones

So far our tractography analysis has revealed converging anatomical projections from orbitofrontal, dorsolateral prefrontal and posterior parietal areas into the striatum. If these do, in fact, reflect an integrative functional network, then cortical areas that show a high degree of anatomical connectivity to the convergent fields should also show significant functional connectivity to these striatal regions. To that end, we used rsfMRI data to measure the functional connectivity between cortical areas and each of the striatal convergence zones. The caudate convergence zones were functionally correlated with a distributed set of bilateral cortical areas, including the DLPFC, lateral OFC, sensorimotor areas, and, most importantly, posterior parietal regions (Figure 5). The connectivity with the parietal cortex appeared to be restricted along the intraparietal sulcus (IPS) and up into the superior parietal lobule (SPL). While convergence zones from both nuclei exhibited a similar pattern of correlated functional activity, several bilateral cortical regions including the pre- and postcentral gyri, lateral occipital and anterior temporal lobes showed functional connectivity with only the putamen convergence zones (Figure 5). However, we also observed a large amount of overlap between the caudate and putamen connectivity to cortex. This overlap was particularly strong in bilateral OFC, DLPFC, and parietal meta-regions. Left and right caudate connectivity appears in more rostral areas of DLPFC and more lateral areas of OFC when compared with putamen connectivity. In parietal cortex, caudate and putamen connectivity overlap to a substantial extent in the IPS, bilaterally, although, there is additional caudate connectivity in posterior regions of both the IPS and the inferior parietal lobule (IPL) that is isolated from putamen connectivity in more anterior IPS.

**Figure 5.**
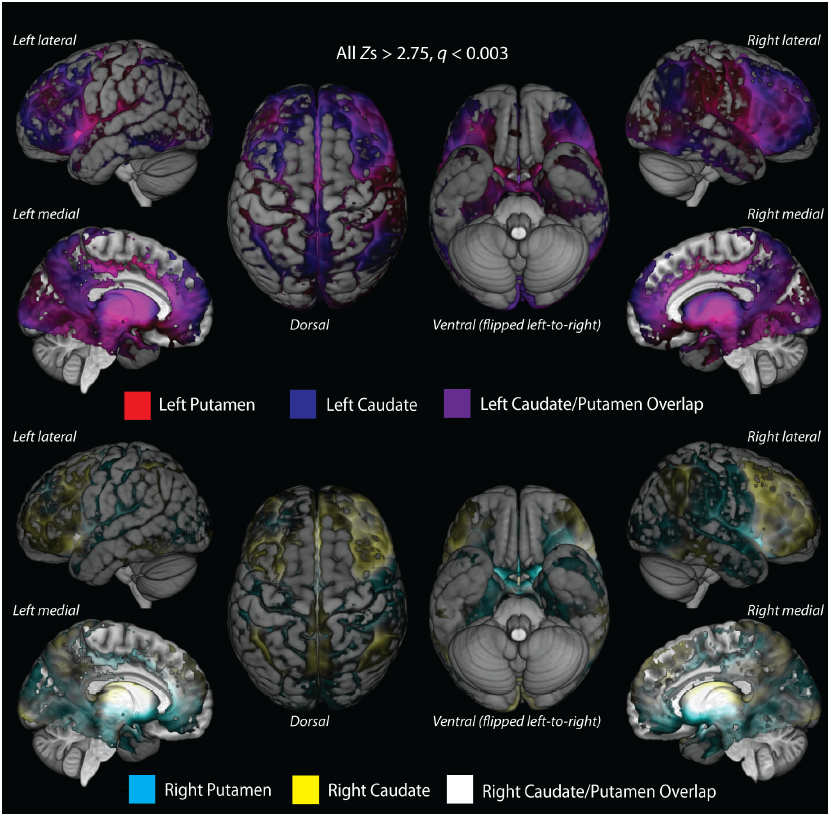
Resting state fMRI (rsfMRI) analysis of cortical regions with significant functional connectivity into each striatal convergence zone, after adjusting for multiple comparisons. Overlaid cortical activity patterns show connectivity to the left (top row of image) and right (bottom row of image) striatum, separately. Cortical surface activity correlated with the left caudate nucleus and putamen convergence zones are represented by blue and red, respectively, while functional connectivity overlap areas are violet. Activation correlated with the right caudate is yellow and right putamen is cyan with white areas of overlapping connectivity.

A visual inspection of Figure 4C and Figure 5 provides qualitative evidence for overlapping structural and functional connections from the striatal convergence zones. In order to quantify this degree of overlap, we calculated the probability that structurally connected voxels were also functionally connected, i.e.,

*P* (*connection*_*fMRI*_ | *connection*_*DSI*_) (see *Methods, Structural and Functional Connectivity Overlap Analysis*) and conducted a bootstrap analysis to construct 95% confidence intervals of mean overlap probabilities. The large percentage of significant structural voxels that also had significant functional connectivity with the convergence zones was consistent across all four striatal convergence zones: left caudate = 66% [24-28%; lower and upper bounds of the 95% chance confidence interval], left putamen = 62% [25-26%], right caudate = 80% [18-30%], and right putamen = 72% [25-27%]. This provides confirmation that voxels showing direct anatomical connections to the striatal convergence zones have a high likelihood—well above chance—of being associated in their functional dynamics.

## Discussion

Our results show that within both the caudate nucleus and the putamen, there are areas that concurrently exhibit structural and functional connectivity to orbitofrontal, dorsolateral prefrontal, and posterior parietal regions of cortex. The location of these convergence zones is anatomically consistent with previous reports of parietal (Selemon & Goldman-Rakic, 1988) and frontal (Haber, Kunishio, Mizobuchi, & Lynd-Balta, 1995; Selemon & Goldman-Rakic, 1985) white matter projections, based on ex-vivo nonhuman primate histology. While the distribution of cortical regions associated with the striatal convergence zones differed to some degree between structural and functional measures of connectivity, a vast majority of cortical areas structurally connected to the convergence zones also showed strong functional connectivity. This supports the notion that these corticostriatal projections form an integrative functional circuit.

Our analysis found that fiber streamlines starting in OFC terminated in ventromedial areas of the rostral striatum, followed caudally by DLPFC projections to more dorsolateral and central striatal regions and, finally, parietal projections had the most dorsolateral and caudal striatal endpoints. This is consistent with a rostral-to-caudal topography of corticostriatal projections that has been previously demonstrated in nonhuman primate histology (Cavada & Goldman-Rakic, 1991; Selemon & Goldman-Rakic, 1985, 1988). Along with this macroscopic topography, at a more local level, histological work has shown that some corticostriatal pathways from disparate cortical areas also have overlapping termination fields in the striatum (Flaherty & Graybiel, 1993; Haber, Kim, Mailly, & Calzavara, 2006; Haber, 2003). Accordingly, we observed clusters of voxels (i.e., convergence zones) bilaterally within both striatal nuclei where projections from several cortical areas including OFC, DLPFC, and posterior parietal cortex terminated. Striatal convergence zone voxels also exhibited significant functional connectivity with structurally connected voxels in the cortex. This is in line with recent work showing that, in humans, distinct striatal regions are functionally connected with networks of distributed cortical areas including the fronto-parietal association, default mode, and limbic networks (Choi *et al*., 2012). While previous work has separately shown projections from OFC (Haber *et al*., 2006; Selemon & Goldman-Rakic, 1985) and posterior parietal cortex (Cavada & Goldman-Rakic, 1991; Choi *et al*., 2012) to overlap with DLPFC projections in the striatum, to the best of our knowledge the present findings show, for the first time, a specific convergence of projections from all three cortical areas to a common striatal area.

Based on previous literature, we propose that this pattern of convergent connectivity may reflect a potential neural mechanism for integrating reward processing, executive control, and spatial attention. In order for reward to bias spatial attention, signals from cortical areas linked to attention must be integrated with reinforcement learning processes, namely, evaluating rewards and using them to shape response selection. Evaluative processes of OFC have been shown to interact with executive control process—e.g., decision-making, goal representation, and action selection (Miller & Cohen, 2001)—to influence goal-directed action selections (O’Doherty, 2004; Hare, O’Doherty, Camerer, Schultz, & Rangel, 2008; Schoenbaum, Roesch, Stalnaker, & Yuji, 2010). Functionally, OFC has been implicated in providing reinforcement signals that influence behavior, with the medial OFC likely representing reward information and lateral OFC representing punishment outcomes (O’Doherty, Critchley, Deichmann, & Dolan, 2003). Additionally, while selective spatial attention processes, linked to the posterior parietal cortex, can influence decision-making (Behrmann, Geng, & Shomstein, 2004; Colby & Goldberg, 1999; Gottlieb, 2007), more recent behavioral research suggests that spatial attention does, in fact, interact with reward as well. Specifically, reward signals may serve to refine behavioral action selections and strategies, reflected by improved efficiency during visual search for highly rewarded spatial targets versus targets that are less rewarded (Della Libera & Chelazzi, 2006; Kristjansson *et al*., 2010; Lee & Shomstein, 2014; Navalpakkam, Koch, Rangel, & Perona, 2010). At the neural level, performance on spatial reinforcement tasks has been shown to be associated with concurrent activity of posterior parietal and DLPFC areas (Lee & Shomstein, 2013a). These independent lines of research provide evidence for the influence of reward and spatial attention on executive control process via functional connections between orbitofrontal and posterior parietal cortices with DLPFC.

Notwithstanding the implications of previous human functional neuroimaging results, it remains unclear whether the integration of these functions is structurally mediated by direct cortico-cortical or interdigitating corticostriatal projections. It is possible that at least part of the interaction between OFC and DLPFC functions is mediated by direct structural connections (Ridderinkhof, van den Wildenberg, Segalowitz, & Carter, 2004); however, our current findings are consistent with a model in which part of this integration may happen at the corticostriatal level (Haber *et al*., 2006). Similarly, histological work supports potential models of spatial attention and executive control integration via direct cortical connections between posterior parietal cortex and DLPFC (Cavada & Goldman-Rakic, 1989), as well as overlapping corticostriatal projections (Cavada & Goldman-Rakic, 1991). While we cannot rule out a direct cortico-cortical connectivity hypothesis, our findings afford some confirmation for the integration of spatial attention and executive control signals in striatal areas that also receive inputs from the OFC, which is consistent with a corticostriatal mechanism for spatial reinforcement learning.

While this integration at the corticostriatal level may seem at odds with the standard view of corticostriatal networks being parallel and independent systems at a global level (see Alexander *et al*., 1986 for review), histological evidence supports the idea of integration at a local level such that disparate cortical areas, within a common circuit, can send sparse projections to the same striatal region (see Haber, 2003 for review). For example, immunohistochemical tracing studies find that cortical projections, from multiple frontal and prefrontal areas, are interdigitated within the striatum of nonhuman primates (Haber *et al*., 2006). Previous diffusion imaging studies in humans have also shown the overlap of corticostriatal projection termination fields in the same striatal voxels (Draganski *et al*., 2008; Verstynen *et al*., 2012). Our identification of bilateral striatal convergence zones is in line with this previous work in both humans and nonhuman primates. Importantly, we observed convergence zones that extended into the dorsomedial caudate nucleus, which has been strongly implicated in reinforcement learning in human functional neuroimaging studies (Badre & Frank, 2012; Daw, Joel, & O’Doherty, 2007; Delgado *et al*., 2005; O’Doherty *et al*., 2004; Schönberg, Daw, Joel, & O’Doherty, 2007). When these previous studies are considered together with our coincidental observation of structural and functional connectivity between OFC, DLPFC, and posterior parietal cortex and the striatum, the convergence of these three corticostriatal pathways, particularly within the dorsomedial caudate, may underlie context-dependent, spatial reinforcement learning suggested in previous research (Lee & Shomstein, 2013b; Nieuwenhuis, Slagter, von Geusau, Heslenfeld, & Holroyd, 2005; Nieuwenhuis, Heslenfeld, *et al*., 2005).

It should be noted that one critical limitation for inferring whether our current results isolate a mechanism for integrating cortical input into the striatum is the relatively low spatial resolution (e.g., 2.4mm isotropic voxels) of current diffusion MRI approaches. We also employed a deterministic tractography method that yields estimates of underlying white matter pathways based on patterns of water diffusion constrained by axonal geometry; however, these streamlines are indirect reflections of the true neuroanatomy (Jones, 2008). Considering these methodological limitations, we cannot directly infer whether the pathways we visualized are actually converging on the same striatal cells or merely terminating in adjacent regions of a nucleus, which can only be verified histologically. Another concern relates generally to rsfMRI functional connectivity analyses, which is an indirect measure of connectivity based on correlated activity throughout the brain. At the time-scale of the BOLD response it is impossible to differentiate direct functional connections to a seed region from indirect connections (Cole, Smith, & Beckmann, 2010). Thus, our inferences based on rsfMRI data can only imply that connected regions represent a functional circuit, but they cannot confirm that correlated areas are directly connected to each other. Finally, neither DSI nor rsfMRI can confirm the task-relevance of the cortical areas that we examined. In order to directly address our hypothesis that this network reflects a neural substrate for spatial reinforcement learning, future work should look at functions of this network during tasks that require the integration of reward, executive control, and spatial attention.

In spite of these limitations, the present findings still provide clear evidence that projections from OFC, DLPFC, and posterior parietal cortex terminate in common striatal regions. While our results are consistent with several independent findings in primate neuroanatomical literature, no previous study has shown the specific convergence of these three corticostriatal pathways in the human brain. Our current findings highlight a plausible structural mechanism that could allow for parietally-mediated spatial attention processes to contribute to the integration of reward and response selection. In future work, the particular dynamics of the neural circuit that we have described here should be explored for their potential role in the integration of spatial attention information with reward and executive control processes during reinforcement learning.

## Acknowledgements

The authors would like to thank Dr. Roberta Klatzky for her consultation regarding data analyses used in this document. This research was sponsored by the PA Department of Health Formula Award #SAP4100062201 and by the Army Research Laboratory under Cooperative Agreement Number W911NF-10-2-0022. The views and conclusions contained in this document are those of the authors and should not be interpreted as representing the official policies, either expressed or implied, of the Army Research Laboratory or the U.S. Government. The U.S. Government is authorized to reproduce and distribute reprints for Government purposes notwithstanding any copyright notation herein.

## Notes

**Conflict of Interest:** The authors do not have any conflicts of interest to report.

